# PD-L1 regulates inflammatory macrophage development from human pluripotent stem cells by maintaining interferon-gamma signal

**DOI:** 10.1101/2022.12.14.520176

**Authors:** Handi Cao, Yang Xiang, Shihui Zhang, Yiming Chao, Jilong Guo, Joshua W. K. Ho, Yuanhua Huang, Pentao Liu, Ryohichi Sugimura

**Affiliations:** Centre for Translational Stem Cell Biology, Hong Kong; School of Biomedical Sciences, Li Ka Shing Faculty of Medicine, The University of Hong Kong; Laboratory of Data Discovery for Health Limited (D24H), Hong Kong Science Park, Hong Kong

**Author notes:** These authors contributed equally (Handi Cao, Yang Xiang, Shihui Zhang).

## Abstract

PD-L1 (programmed death-ligand 1) serves as a pivotal immune checkpoint in both the innate and adaptive immune systems. PD-L1 is expressed in macrophages in response to interferon-gamma (IFNγ). We examined whether PD-L1 might regulate macrophage development. We established *PD-L1*^*-/-*^ human pluripotent stem cells, differentiated them into macrophages, and observed a 60% reduction of CD11B^+^CD45^+^ macrophages in *PD-L1*^*-/-*^, orthogonally verified with PD-L1 inhibitor BMS-1166 reduced macrophages to the same fold. Single-cell RNA sequencing further confirmed the 60% reduction of macrophages as well as the down-regulation of macrophage-defining transcription factors *SPI1, KLF6*, and *MAFB*. Further, *PD-L1*^*-/-*^ macrophages reduced the level of inflammatory signals such as NFκB, TNF, and chemokines CXCL and CCL families. Whilst anti-inflammatory TGF-β was upregulated. Finally, we identified that *PD-L1*^*-/-*^ macrophages significantly down-regulated interferon-stimulated genes (ISGs) despite IFNγ in differentiation media. Mechanistically, *PD-L1*^*-/-*^ macrophages reduced *IFNGR1* expression explaining that cells could not respond to IFNγ. These data suggest that PD-L1 regulates inflammatory macrophage development by maintaining the IFNγ signal.

## Introduction

Programmed death ligand 1 (PD-L1/CD274/B7-H1) is abundant in the tumor and suppresses T-cells (*1, 2*). PD-L1, expressed in cancer or myeloid cells, binds with its receptor PD-1 in T-cells (*3*). The phosphorylated cytoplasmic domain of PD-1 recruits SHP-2, resulting in the down-regulation of T-cell signaling pathways (*4, 5*). PD-L1 is one of the major targets of cancer immunotherapy (*6*). Its blockade results in reactivation and clonal-proliferation of antigen-experienced T cells in the tumor (*7*). Whether PD-L1 blocking therapy is successful depends on the tumor’s immunogenicity (*8*). For example, interferon-gamma (IFNγ) restricts tumor development, however, continuous exposure results in immune escape (*9*). Durvalumab, a fully-human IgG1 antibody against PD-L1, was FDA-approved and demonstrated progression-free survival significantly longer than with a placebo in Stage III Non–Small-Cell Lung Cancer (*10*). In contrast, Avelumab, another fully-human IgG1 antibody against PD-L1 failed phase III clinical trials for gastric cancer (*11*). These clinical trials suggest the significance of combination therapy targeting immune checkpoint therapy resistance mechanisms (*12*).

Interferon-gamma (IFNγ) secreted in the tumor microenvironment promotes PD-L1 expressions on both cancer cells and myeloid cells (*13*). IFNγ is predominantly expressed by Th1-cells, cytotoxic T-cells, and NK-cells in the tumor (*14*). IFNγ binds with its receptor IFNGR1 which phosphorylates STAT1. The dimerized phospho-STAT1 binds to promoters of interferon-stimulated genes (ISGs) (*15*). IRF1, one of the ISGs, is a transcription factor that induces PD-L1 expression (*13*). Cells expressing IFNGR1 can respond to IFNγ. Major responders in the tumor are cancer cells, myeloid cells including macrophages, regulatory T-cells, Th1-cells, cytotoxic T-cells, and NK cells (*16*). Myeloid cells are the predominant immune cell population that responds to IFNγ and express PD-L1 in the tumor (*17*). IFNγ-induced expression of PD-L1 is considered a feedback mechanism, resulting in PD-L1 on cancer cells and myeloid cells that block T-cells (*18*). However, the low expression level of intratumoral IFNγ in invasive cervical carcinomas correlated with poor clinical prognosis (*19*). Administration of recombinant IFNγ enhances the efficacy of CAR-T cell therapy (*20, 21*), consistently that IFNγ works in favor of anti-cancer immunity, whilst inducing PD-L1. High expression of PD-L1 in the tumor correlated with better prognosis in non-small lung carcinoma (*22*). Meta-analysis of breast cancer patients revealed that high expression of PD-L1 in the tumor correlated with better prognosis (*23*). These indicate a potential mechanism that PD-L1 supports anti-cancer immunity in a certain context of cancers.

PD-L1 is a type I transmembrane glycoprotein with a short intracellular domain. Although the major role of PD-L1 had been considered merely binding with its receptor PD-1, the intracellular domain of PD-L1 facilitates signal transduction cell-autonomously (*24, 25*). PD-L1 directly prevents cancer cell apoptosis by inhibiting cytotoxic activities from type I and type II IFN (*24*) and promoting cancer cell proliferation via mTOR signal (*26*). Nuclear PD-L1 functions as a transcription factor and induces necrosis and immune-response genes in cancer cells (*27, 28*). These indicate that PD-L1 facilitates signal transduction independent from its receptor PD-1.

The role of PD-L1 is investigated in the PD-1 T-cells context predominantly (*29*). The knockout of *PD-L1* results in the activation of cytotoxic T-cells and the generation of the pro-inflammatory milieu in tissues (*30*). However, a few studies suggested the role of PD-L1 in immune cells independent from the PD-1 T-cells context. PD-L1 marks pre-hematopoietic stem cells in mouse embryos where IFNγ plays a key role in the emergence of these cells (*31*–*33*). The knockout of PD-L1 in adult macrophages, both from humans and mice, resulted in the activation of inflammatory macrophages, suggesting that PD-L1 may suppress inflammatory programs in adult tissue-derived macrophages (*34*).

IFNγ is a major inducer of inflammatory macrophages (*35*). IFNγ signal increases responsiveness to pro-inflammatory stimuli such as lipopolysaccharide (LPS) or type I interferons and resistance to anti-inflammatory stimuli such as IL-4, IL-10, and glucocorticoids (*36*). Hence IFNγ activates inflammatory programs. Meanwhile, IFNγ induces PD-L1 expression in cancer cells to prevent necrosis (*13, 24*), whilst the PD-L1 role in macrophages is considered to suppress inflammatory programs. Macrophages, a versatile cell source for immunotherapy (*37*–*39*), and their fine-tuning of inflammatory programs are the key to successful cell therapy against cancer and fibrosis (*40*).

Here we modeled human macrophage development in yolk sac organoids and examined the role of PD-L1. Both genetic and pharmacological inhibition of PD-L1 impaired macrophage development. The resultant macrophages defected the inflammatory program, unable to respond to IFNγ due to the reduction of its receptor IFNGR1. These observations indicate that PD-L1 induced by IFNγ maintains the IFNγ signal and assures the development of inflammatory macrophages. Our study demonstrates that PD-L1 regulates IFNγ signal and establishes inflammatory programs of human macrophages.

## Results

We determined the role of PD-L1 in macrophage development from human pluripotent stem cells to yolk sac organoids (Figures 1A and 1B). We formed EBs (embryoid bodies) and applied STEMdiff™ hematopoietic kit for mesoderm patterning and differentiation into yolk sac organoids (*41*). We instructed them into monocytes by STEMdiff™ monocyte differentiation supplement then subsequent maturation to macrophages by ImmunoCult™-SF macrophage medium. We defined macrophages by the expression of CD11B, CD14, CD16, CD45, and CD68 (Figure 1C, Supplementary Figure 1). We polarized macrophages to pro-inflammatory status by LPS and IFNγ, while polarized to anti-inflammatory status by IL-4, respectively. PD-L1 was predominantly expressed in pro-inflammatory macrophages. In contrast, its receptor PD-1 expression could not be detected during the differentiation of yolk sac organoids (Figure 1C). Moreover, PD-L1 came up exclusively upon exposure to IFNγ (Figure 1D).

**Figure 1.**
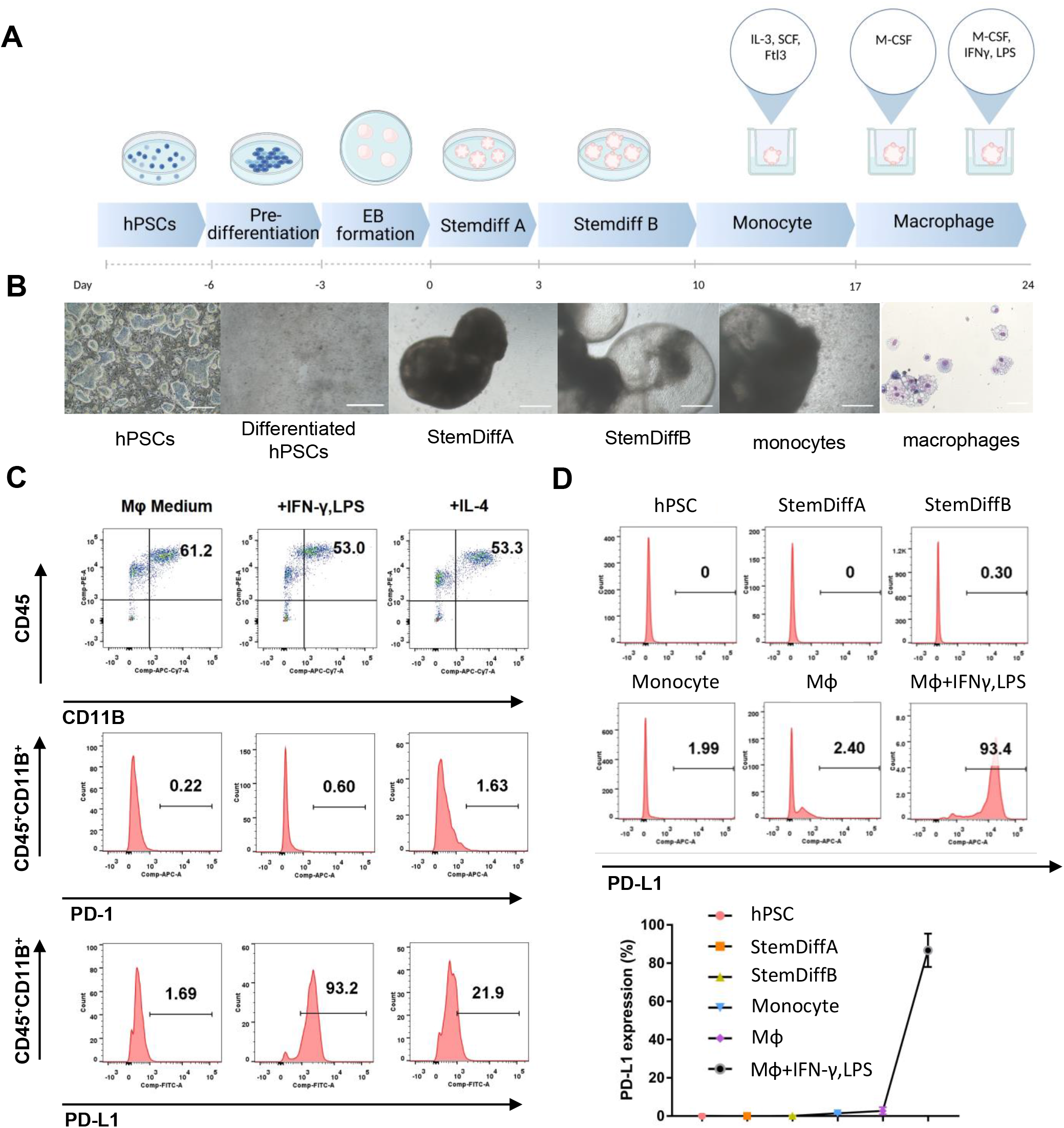
Human macrophages from yolk sac organoids express PD-L1 upon IFNγ stimulation. **A**. Schematic overview of the generation human yolk sac organoids and macrophages. Human pluripotent stem cells (hPSCs) were differentiated into yolk sac organoids by commercial media (StemDiff), then induced macrophages with M-CSF. **B**. Bright-field images of representative cellular morphology from hPSC differentiation into human yolk sac organoids and macrophages. EB: embryoid body, HPC: hematopoietic progenitor cell. **C**. Flow cytometry analysis of CD45, CD11B, PD-1, and PD-L1 on macrophages in macrophage basal medium (Mφ medium), Mφ medium plus IFNγ, and LPS, Mφ medium plus IL-4, respectively. **D**. Flow cytometry analysis of PD-L1 expression in hPSCs, human yolk sac organoids (StemDiffA and StemDiffB), and macrophages. The representative data of flow cytometry were shown here.

To examine the role of PD-L1 in macrophage development, we established *PD-L1*^*-/-*^ human pluripotent stem cells by targeting exon 2 of the *CD274* gene which encodes PD-L1 (Supplementary Figure 2). We confirmed knockout efficiency (Figures 2A, B). The percentage of CD45^+^CD11B^+^ macrophages significantly decreased in *PD-L1*^*-/-*^ yolk sac organoids, suggesting that PD-L1 regulates macrophage development (Figures 2C, D). Two *independent PD-L1*^*-/-*^ human pluripotent stem cell lines A7-1 and B2-3 showed a similar reduction of macrophages (Supplementary Figure 3). PD-L1 inhibitor BMS-1166 (*42*) phenocopied the knockout lines by reducing the percentage of CD45^+^CD11B^+^ macrophages (Figure 2E). These data suggest that PD-L1 supports the development of macrophages in yolk sac organoids.

**Figure 2.**
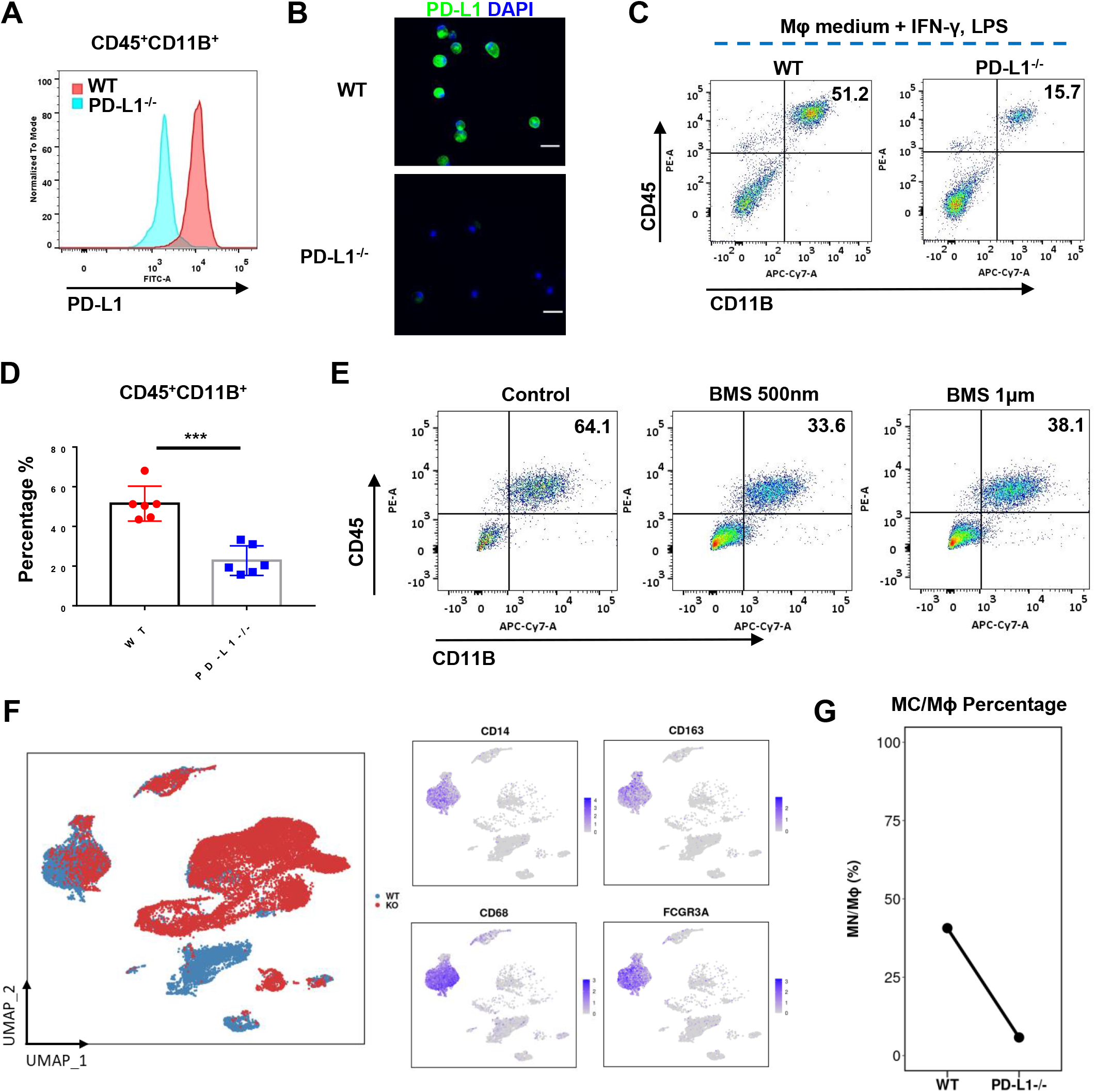
Knockout of *PD-L1* prevents macrophage development. **A-B**. Flow cytometry analysis (A) and immunofluorescence (B) of PD-L1 expression on *wild type* and *PD-L1*^*-/-*^ macrophages in Mφ medium plus IFNγ and LPS. Scale bar: 50 μm. **C-D**. Flow cytometry analysis of macrophage (CD45^+^CD11B^+^) (C) and percentage (D) of *wild type* and *PD-L1*^*-/-*^ macrophages in Mφ medium plus IFNγ and LPS. Statistical results shown here were from three independent experiments (n=6). ***, *p*<0.001. **E**. PD-L1 inhibitor BMS-1166 reduced macrophage development in human yolk sac organoids. Schematic overview of BMS-1166 treatment timeline on macrophage differentiation in the human yolk sac (YS) organoids (upper). Flow cytometry analysis of CD45, CD11B on macrophages after 500nm or 1μm BMS-1166 treatment (below). The representative data of flow cytometry were shown here. **F**. Uniform manifold approximation and projection (UMAP) visualization of single cells from yolk sac organoid in *wild type* (WT, *Num. of Cells = 9181*, color in blue) and *PD-L1* knock-out (KO, *Num. of Cells = 16258*, color in red). Each dot represents a single cell. In total 19 cell types were annotated and displayed in different colors (See Supplementary Figure 5). The right panel shows the expression of macrophages and monocyte-specific marker genes. **G**. Dot-line chart displaying the decrease of monocyte/macrophages (MC/Mϕ) proportion upon *PD-L1* knockout. Statistical significance (p-value) is calculated by the likelihood ratio test and adjusted by Benjamini & Hochberg method by R package DCATS.

To further characterize *PD-L1*^*-/-*^ macrophages, we performed 10X Chromium single-cell RNA-sequencing (scRNA-seq). We sequenced whole-live cells from yolk sac organoids after induction to pro-inflammatory macrophages. We subsequently removed low-quality cells and doublets with DoubletFinder (Supplementary Figure 4). We obtained a total of 25,439 cells, with 16,258 from *wild type* and 9,181 from *PD-L1*^*-/-*^ yolk sac organoids respectively. We annotated 19 cell types according to canonical markers and visualized them by UMAP. We defined the monocytes and macrophages (MC/Mφ) population expressing *CD14, CD68, CD163*, and *FCGR3A* (encodes CD16) (Figure 2F, Supplementary Figure 4). We consistently detected a significant reduction of macrophages in both flow cytometry and scRNA-seq (Figure 2D vs. Figure 2G).

We next determined the differentially expressed genes (DEGs) in MC/Mφ between *WT* and *PD-L1*^*-/-*^ yolk sac organoids (Figure 3A; Supplementary Figure 5). *PD-L1*^*-/-*^ MC/Mφ reduced the expression of macrophage-determining transcription factors (TFs) *SPI1, KLF6*, and *MAFB* (Figure 3B). The Gene Ontology (GO) and KEGG analysis by GSEA revealed the significant down-regulation of pro-inflammatory associated signaling pathways, such as NOD, chemokine, TNF, NF-kB, IL-17, antigen presentation signal pathways, in contrast, up-regulation of an anti-inflammatory TGF-β signaling pathway (Figure 3C). This suggests that PD-L1 may regulate inflammatory macrophage development.

**Figure 3.**
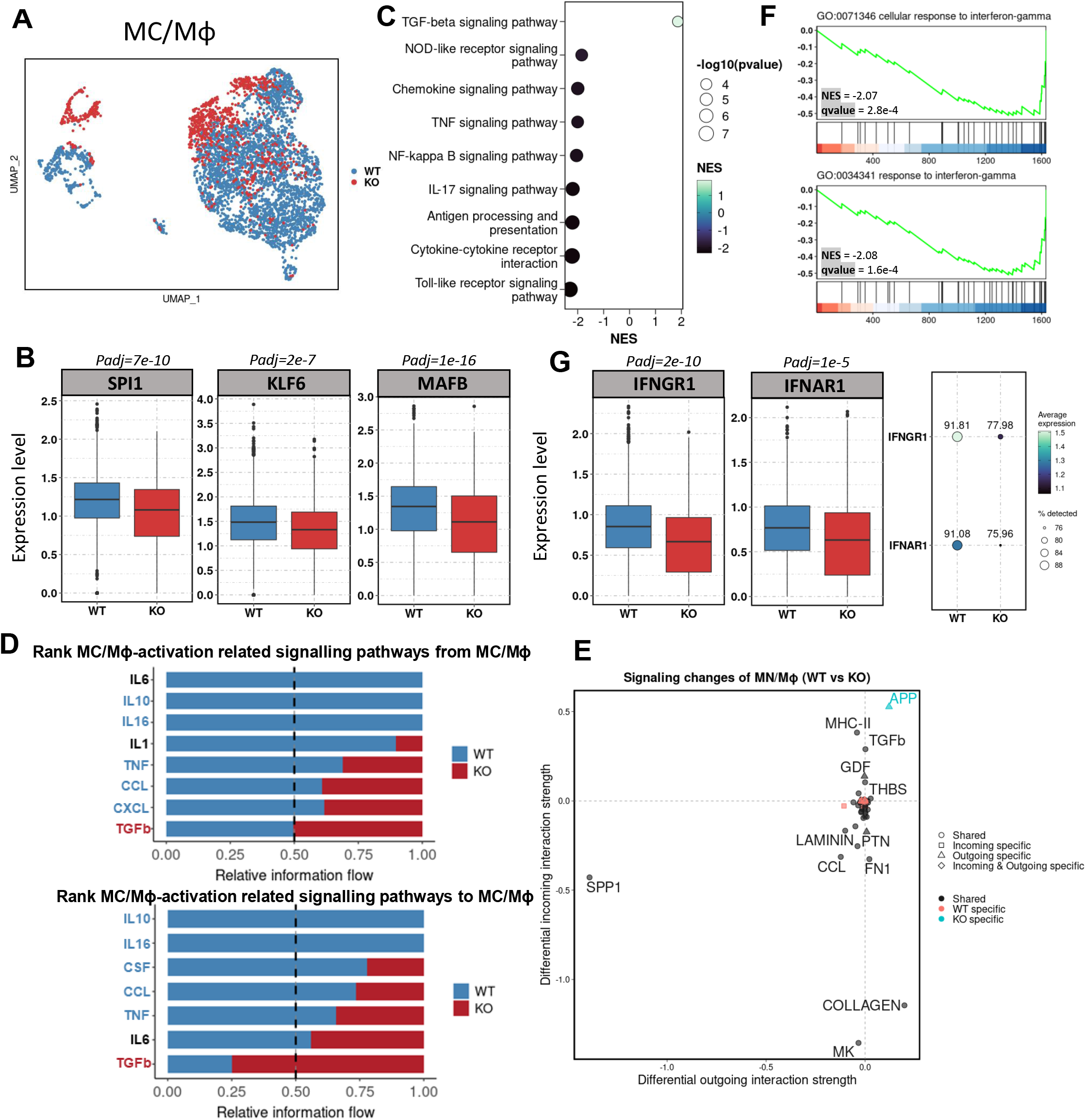
*PD-L1*^*-/-*^ macrophages down-regulate inflammatory programs and are unable to respond to IFNγ. **A**. UMAP visualization of monocyte and macrophages (MC/Mϕ) population, colored by the genotype. (WT: wild-type; KO: *PD-L1* knockout) **B**. Box-plot showing the decreased expression of selected macrophage-determining transcription factor within the MC/Mϕ cluster upon *PD-L1* knockout. (p-value calculated by “bimod” test in Seurat R package, corrected by “Benjamini-Hochberg” method). **C**. Selected KEGG pathways enriched by gene set enrichment analysis (GSEA) of differentially expressed genes (DEGs) between *PD-L1* knockout (KO) and wild-type (WT) within the MC/Mϕ cluster. NES, normalized enrichment score. DEGs are calculated using the same method in Figure 3B. **D**. Bar graphs showing the ranking of major outgoing (up-panel) and incoming(down-panel) signals of MC/Mϕ upon *PD-L1* knockout (KO) compared to wild-type (WT), respectively. The rank of signals was based on differences in the overall information flow of each group. **E**. Scatter-plot visualizes the differential outgoing and incoming signaling associated with monocyte/macrophages group between *WT* and *PD-L1 KO* group. Positive values indicate the increase in the KO group, while negative values indicate the increase in the *WT* dataset. Analysis by *CellChat* R package. **F**. GSEA analysis shows the enrichment of differentially expressed down-regulated genes upon *PD-L1* knockout in MC/Mϕ cells into interferon-gamma response. **G**. Box-plot and Dot-plot showing the reduction of both expression and percentage of expressing cells of interferon-gamma receptor *IFNGR1* and interferon-alpha receptor *IFNAR1* within MC/Mϕ cluster upon *PD-L1* knockout, respectively. (p-value calculated by “bimod” test in Seurat R package, corrected by “Benjamini-Hochberg” method).

Consistently, cell-cell communication analysis showed the reduced macrophage activation ligand-receptor network in the *PD-L1*^*-/-*^ MC/Mφ such as IL6, IL10, CSF, and chemokines CCL and CXCL. Whilst, *PD-L1*^*-/-*^ MC/Mφ increased the anti-inflammatory TGF-β ligand-receptor network (Figure 3D). We identified GDF and THBS signals increased (Figure 3E). GDF15 called macrophage-inhibitory cytokine-1 (*43*) significantly interacted with TGFBR2. THBS1 interacted with macrophage suppressor CD47 (*44*) (Supplementary Figure 6). These data indicate that *PD-L1*^*-/-*^ MC/Mφ were anti-inflammatory.

We then examined if *PD-L1*^*-/-*^ MC/Mφ responds to IFNγ present in macrophage activation media (Figure 1A). GSEA analysis revealed that interferon-stimulated genes (ISGs) were significantly down-regulated in *PD-L1*^*-/-*^ MC/Mφ, suggesting that these cells could not respond to IFNγ (Figure 3F). *PD-L1*^*-/-*^ MC/Mφ reduced the expression of interferon receptors IFNGR1 and IFNAR1 (Figure 3G). These results demonstrate that PD-L1 may maintain the IFNγ signal by sustaining the expression of its receptor IFNGR1 and establishing inflammatory programs.

Finally, we examined the resemblance of human macrophages between yolk sac organoids and fetal yolk sacs (*45*). We identified that cellular components such as macrophages (MC/Mφ), common myeloid progenitors (CMP), granulocytes, and yolk-sac myeloid-biased progenitors (YSMP) were correlated between yolk sac organoids (labeled as PD-L1 WT) and fetal yolk sac (labeled as In Vivo YS) (Supplementary Figure 7). Taken together, our data indicate that PD-L1 regulates the development of inflammatory macrophages in human yolk sacs. Mechanistically, PD-L1 maintains the IFNγ signal to establish inflammatory programs in macrophages.

## Discussion

PD-L1 inhibition reduced inflammatory macrophages as well as ISGs and IFNGR1 expression, resulting that *PD-L1* ^*-/-*^ macrophages could not respond to IFNγ. Our data suggest that PD-L1 maintains the integrity of the IFNγ signal by sustaining the expression of IFNGR1. How and whether PD-L1 directly regulates ISGs and IFNGR1 expression is intriguing. The reduction of macrophage-determining TFs by *PD-L1* knockout implies that PD-L1 governs the gene-regulatory network of macrophage development. The nuclear PD-L1 might be a potential mechanism to maintain the IFNγ signal and macrophage gene-regulatory network (*27, 28*).

The use of human pluripotent stem cells (hPSCs) offered us the advantage of studying human macrophage development in the yolk sac. We previously established yolk sac organoids from hPSCs that commit to erythro-megakaryocyte and monocyte lineages (*41*). In this study, we employed M-CSF to induce macrophages in yolk sac organoids. Macrophages in our study resemble those of fetal stage yolk sacs as indicated by cross-reference of scRNA-sequencing datasets (*45*). The phenotype reported in this study is independent of T-cells, whose over-activation usually masks intrinsic changes in *PD-L1*^*-/-*^ macrophages (*30*). PD-L1 in yolk sac macrophages may regulate the establishment of inflammatory programs by maintaining the IFNγ signal. This would explain the difference observed in another study where PD-L1 was immune suppressive in adult macrophages that are of bone marrow origin (*34*). Macrophages in the tumors are initially of yolk sac origin that supports anti-cancer immune cells, then migrated by those of bone marrow origin, the latter promoting cancer growth (*46, 47*). Our study proposes that PD-L1 establishes an inflammatory program in yolk sac-origin macrophages. Consistently, the phase II trial of PD-L1 inhibitor durvalumab in patients with advanced esophageal and gastroesophageal junction adenocarcinoma (NCT02639065) increased the proportion of anti-inflammatory macrophages in the tumor, a possible adverse effect on yolk sac origin macrophages (*48*). Our results suggest the potential clinical significance of the increase of anti-inflammatory macrophages in PD-L1 inhibitor treatment.

## Methods

### Generation of macrophages from human pluripotent stem cells

#### 1. Human pluripotent stem cell (hPSC) maintenance and pre-differentiation

hPSCs were obtained from Pentao Liu’s lab and maintained in the medium with chemically defined conditions (*49*). Cells were first pre-differentiated in KSR medium (DMEM/F12+10% KSR) for 2 days then followed by embryoid body (EB) formation.

#### 2. EB formation

Remove the KSR medium and wash the cells with PBS. Digest cells with TrypLE™ (12605036, Gibco) at 37 °C for around 5 minutes. Remove TrypLE™ and add KSR medium to harvest cells. Avoid harsh pipetting. Centrifuge cells under 300g for 3 minutes. Remove the supernatant carefully. Add KSR medium and ROCK inhibitor Y-27632 (10μM) to resuspend the cells. Each 30μL hanging drop containing 5000 cells was made onto the inside of the lid of a 100mm petri dish (20100, SPL). Make 30-40 drops for each lid and fill the dish with PBS to keep moist. Invert the lid gently to cover the dish. Incubate the EBs for 3 days.

#### 3. Macrophage generation from yolk sac organoids

Collect all EBs with a 1000uL pipette and centrifuge them under 20g for 1 min. Remove the supernatant carefully. Culture EBs in an ultra-low-adherence plate with Stemdiff A (STEMdiff™ Hematopoietic Kit, Stem cell technologies) for 3 days and Stemdiff B (STEMdiff™ Hematopoietic Kit, Stem cell technologies) for 7 days as described previously (*41*). Then all cell clumps formed in Stemdiff B were transferred to a 24-well transwell (0.4uM pore, Stem cell technologies, #38024) for further differentiation. Monocyte induction medium (StemSpan™ SFEM II plus monocyte differentiation supplement) was used for monocyte generation and then macrophage induction medium (ImmunoCult™-SF Macrophage Medium containing 50ng/mL M-CSF) was applied for macrophage generation. Macrophage media was supplied with M-CSF, IFN-γ, and LPS.

### Establishment of PD-L1^-/-^ hPSCs by CRISPR-Cas9

sgRNAs were designed to target the *CD274/PD-L1* exon2. The designs were referred to https://chopchop.cbu.uib.no/. sgRNAs were synthesized by IDT technologies. The editing efficiency of these sgRNAs was initially validated on hPSCs by electroporation. The high-editing sgRNA sequence was shown in Supplementary Figure 2.

The CRISPR-Cas9 knockout was conducted as follows. The synthetic oligos were annealed, then the annealed double-strand oligos and backbone vectors were digested by restriction enzyme BbsI (Thermo Scientific™, catalog no. ER1011). After enzyme digestion, purified the products respectively. Purified oligos and vectors were ligated overnight. On the following day, the bacterial transformation was performed for recombinant PD-L1 KO construct amplification. Then, extracted plasmid sequences were confirmed by Sanger sequencing.

*PD-L1 KO* piggyBac plasmids were generously provided by Pentao Liu’s lab. hPSCs were maintained for 4-5 days before electroporation. The confluency was around 70%∼80%. We used the Human Stem Cell Nucleofector Kit 2 (Lonza, catalog no. LONVPH-5022). After electroporation, hPSCs were re-seeded on the plate and the medium should be changed the following day. Three days later, cells were selected by the addition of 2 ug/mL puromycin (Sigma-Aldrich, catalog no. P9620-10ML). After several days of selection, the colonies were picked, expanded, and genotyped, respectively. The two pairs of genotyping primers were designed as follows: CD274-1F: AACCGACCAGATAAAGTGATT, CD274-1R: ATCCTGCAATTTCACATCTGTGA; CD274-2F: ATAAACGCTGTGCCAATTTTGT, CD274-2R: TCATGCAGCGGTACACCCCTG. Single colonies confirmed to inherit the deletion by Sanger sequencing were selected for the following experiments.

### Flow cytometry analysis

Cell suspensions were prepared before flow cytometry analysis. For the hPSC line, TrypLE™ was used for dissociation; For EBs and other cell clumps from Stemdiff A & B, monocyte induction medium and macrophage induction medium, ACCUMAX™ (catalog number) was used for dissociation; To collect cells attached in transwell, ACCUMAX™ was also used. Resuspend cells in 100ul FACS staining buffer and add 5ul FcX (human, Biolegend 422302) into each sample at room temperature for 10min. Stain cell surface markers with optimal concentrations of specific antibodies (CD11B-APC-Cy7, CD45-PE, CD14-FITC, CD16-PE-Cy7, CD68-PE-Cy7, PD-1-FITC, PD-L1-APC) for extracellular staining following previous panel design for 30min on ice. Wash cells with PBS once. Centrifuge samples at 400-600g at RT for 5min, and discard the supernatant. DAPI was added to all samples before analysis. Analysis was performed on BD FACSymphony™ A3. For all analyses, single cells were gated based on FSC-A versus FSC-H, and DAPI^+^ dead cells were gated out.

### May-Grunwald-Giemsa staining

All cells after the macrophage stage were collected and used for cytospin on glass slides with the program 800rpm, 3min. After drying slides overnight, slides were immersed in 100% May-Grunwald for 4min followed by washing, then transferred to 4% Giemsa for another 4min. Slides were rinsed in tap water to make sure the background blue staining has been removed. Slides were dried before taking photos under the microscope.

### Immunofluorescence

Fixed WT or KO total cells respectively in 4% formaldehyde (Beyotime, catalog no. P0099-100ml) for 20 minutes at room temperature, then spined down the cells, poured off the supernatant and resuspended the cell pellets in 200 ul cell suspension buffer. After cytospin, then the slides were permeabilized with 0.5% Triton X-100 (Sigma-Aldrich, catalog no. 93443-100ML) for 30 minutes and blocked with 10% BSA solution (Millipore Sigma, catalog no. 126615-25ML) for 1 hour. Washed the slides twice and then stained the slides with the primary antibody anti-PD-L1 (Abcam, catalog no. ab213524) overnight in a wet box. After 12 hours, rinsed the slides thrice with PBS, then added the secondary antibody Goat Anti-Rabbit IgG H&L (Alexa Fluor® 488) (Abcam, catalog no. ab150077) and incubated for 1 hour. Rinsed the slides thrice with PBS and added 100 ul diluted DAPI buffer (BD Bioscience, catalog no. 564907) to each slide, and incubated for 3 minutes at room temperature. Washed the slides twice with PBS and then kept the slides wet. Images were taken by Nikon Ti2E Fluorescent microscope and merged using ImageJ software.

### Single-cell RNA-sequencing

#### 1. Sample preparation

The *wild type* and *PD-L1*^*-/-*^ hPSC lines followed the macrophage differentiation. All cells were collected and resuspended in a FACS buffer. Cells were stained by CD11b-APC-Cy7 and CD45-PE. After staining, cells were washed and resuspended in FACS buffer with DAPI. Cells were sorted using a Fusion Flow Cytometer (BD Bioscience). DAPI-negative cells were sorted for Single-cell RNA-seq with the support of the Centre for PanorOmic Sciences (CPOS). Briefly, the single-cell RNA-seq library was constructed and performed by Chromium Next GEM Single Cell 3ʹ Reagent Kit v3.1. Illumina Novaseq 6000 for Pair-End sequencing services was used and the final data output was 206.2 for *PD-L1*^*-/-*^ and 213.9G for *wild type*.

#### 2. Data analysis

Single-cell transcriptomic sequencing data of *wild-type* and *PD-L1*^*-/-*^ samples from 10X Genomics were processed with CellRanger software (v7.0.0) and mapped to the GRCh38 human reference genome. The read counts of each gene by cell were counted and loaded by the Seurat R package (v4.2.0). The quality control and cell filtering were conducted for two datasets separately. In detail, cells with fewer than 500 detected features, 2000 counts, and more than 15% of mitochondrial gene expression were removed. And doublets detected by the Doublet Finder R package (v2.0.3) with either high or low confidence were subsequently all filtered out.

After quality control, we checked the batch effect between our two datasets using Seurat’s built-in FindIntegrationAnchors function. The top 2,000 highly variable genes were used for principal component analysis (PCA) to conduct the dimension reduction and clustering and further visualized by Uniform Manifold Approximation and Projection (UMAP). Markers of each cluster were identified by the Seurat FindAllMarkers function, which together with well-known canonical gene markers, were used for cell type annotation.

The differentially expressed gene (DEG) analysis on macrophage population (MC/Mφ) between *wild-type* and *PD-L1*^*-/-*^ samples was performed using the Seurat FindMarkers function, with the “bimod” test used. (Likelihood-ratio test for single-cell gene expression). The DEG was defined as the genes with q value < 0.05 and absolute value of average log2 fold change > 0.1.

The gene ontology (GO) and KEGG analysis to the whole DEG result (all genes kept but q value equal to NA) was performed by gene set enrichment analysis (GSEA) using the gseGO and gseKEGG function of clusterProfiler R package (v4.4.4). The enriched GO and KEGG terms of interest, defined by q value < 0.05, absolute value of normalized enrichment score (NES) > 1, were visualized by dot plot.

Cell-cell communication analysis was performed by CellChat R package (v1.5.0). The filtered gene expression raw data of our two datasets with annotated cell type information was loaded to establish the CellChat object respectively. The ligand and receptors (L-R) within different cell populations were then recognized and calculated for L-R pair probability. To compare the cell interactions between *wild-type* and *PD-L1*^*-/-*^ samples, we then merged these two CellChat objects into one. The total number of interactions and strength of cell-cell communication networks were inferred by the compareInteractions function. The information flow of a specific signaling network was compared between *wild-type* and *PD-L1*^*-/-*^ samples and then visualized by rankNet function as a stacked bar plot, with sources and targets set as macrophages group (MC/Mφ) separately.

The statistical significance analysis of differential cell-type abundance between *wild-type* and *PD-L1*^*-/-*^ samples was performed by the DCATS R package (*50*), with parameters by default. The LRT_pval (p-value calculated by likelihood ratio test) less than 0.01 was used to define whether a cell type show differential proportion upon *PD-L1* knockout.

### Statistical analysis

All results were derived from at least three independent experiments. Values are expressed as mean ± SD. Comparison between groups was performed using Student’s unpaired t-test or one-way ANOVA (GraphPad Prism 7.00 Software, Inc., San Diego, CA, USA). P values less than 0.05 were accepted to indicate statistically significant differences.

## Data availability

The scRNA-seq data reported in this study have been deposited in NCBI with the accession number GSE218722. The link to the dataset https://www.ncbi.nlm.nih.gov/geo/query/acc.cgi?acc=GSE218722.

## Acknowledgment

We are thankful to CPOS at HKUMed for technical assistance in scRNA-seq, and FACS. We are thankful to Cheryl Tam for operating the cell sorting. We appreciate American Journal Experts AJE for English proof service. We appreciate Danny Chan, Yuanhua Huang’s lab members for discussion, and members of the Centre for Translational Stem Cell Biology for discussion and administrative support. The study is supported by the Platform for Technology Fund, RGC ECS 27109921, Seed Fund from the School of Biomedical Sciences, and InnoHK Centre for Translational Stem Cell Biology.

## Author contributions

H.C., Y.X., and R.S. conceived the study. H.C. and Y.X. conducted experiments. S.Z. conducted informatics analysis with the guidance of Y.C., Y.H., and R.S. J.H. contributed R package DCATS. Y.X. and J.G. generated *PD-L1* knock-out cells with the guidance of P.T. and R.S. H.C.,Y.X., S.Z., and R.S. wrote the manuscript.

## Figure Legends

**Supplementary Figure 1.**
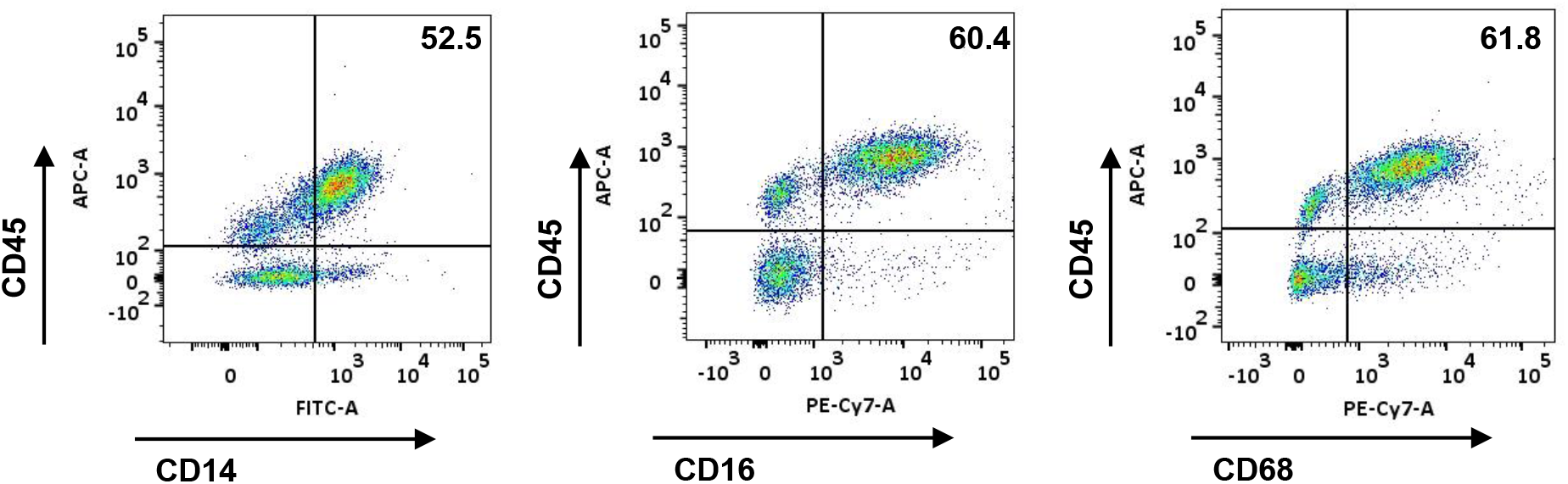
Macrophages from human yolk sac organoids. Flow cytometry analysis of CD14, CD16, CD45, and CD68 on macrophages from human yolk sac organoids. The representative data of flow cytometry were shown here.

**Supplementary Figure 2.**
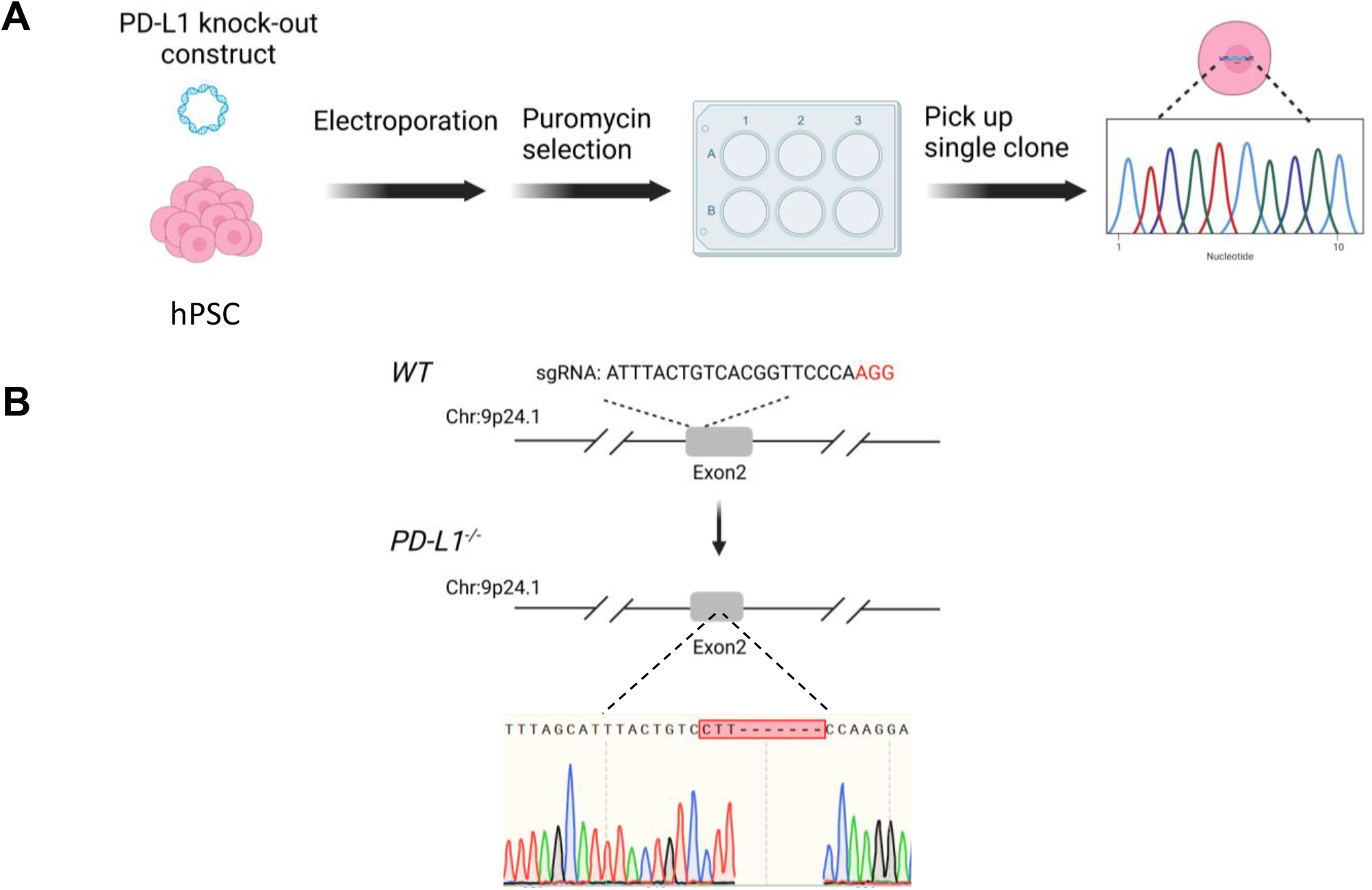
Establishment of *PD-L1*^*-/-*^ human pluripotent stem cell lines. **A**. The PD-L1 genetic knock-out protocol in the EPSC stage. **B**. The genetic knock-out strategy. The sgRNA was designed on WT (wild-type) exon2 (upper panel), and the *PD-L1 (CD274)* locus after targeting was shown (lower panel). Sanger sequencing result reveals the 4 base-pair deletions on KO (knock-out) exon2.

**Supplementary Figure 3.**
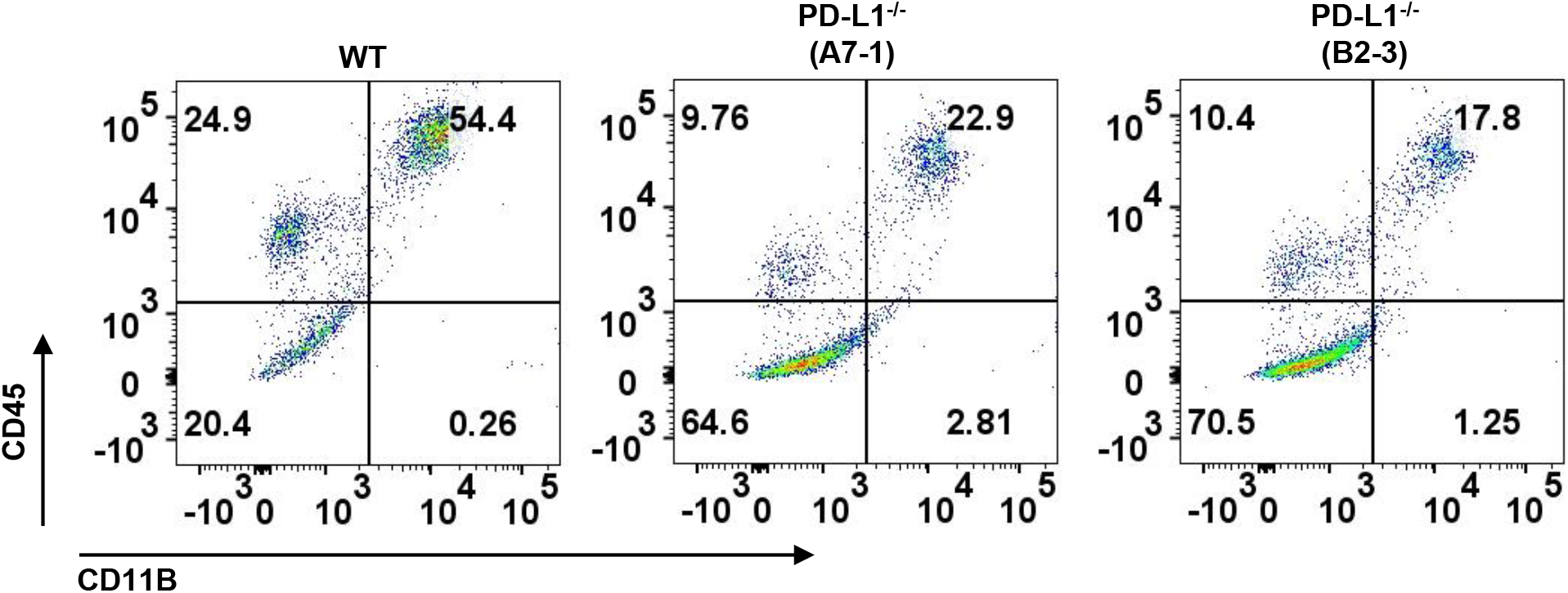
Reduction of macrophage development in independent *PD-L1* knockout lines. Flow cytometry analysis of CD45+CD11B+ macrophages between Wild type vs. KO A7-1 and KO B2-3 lines.

**Supplementary Figure 4.**
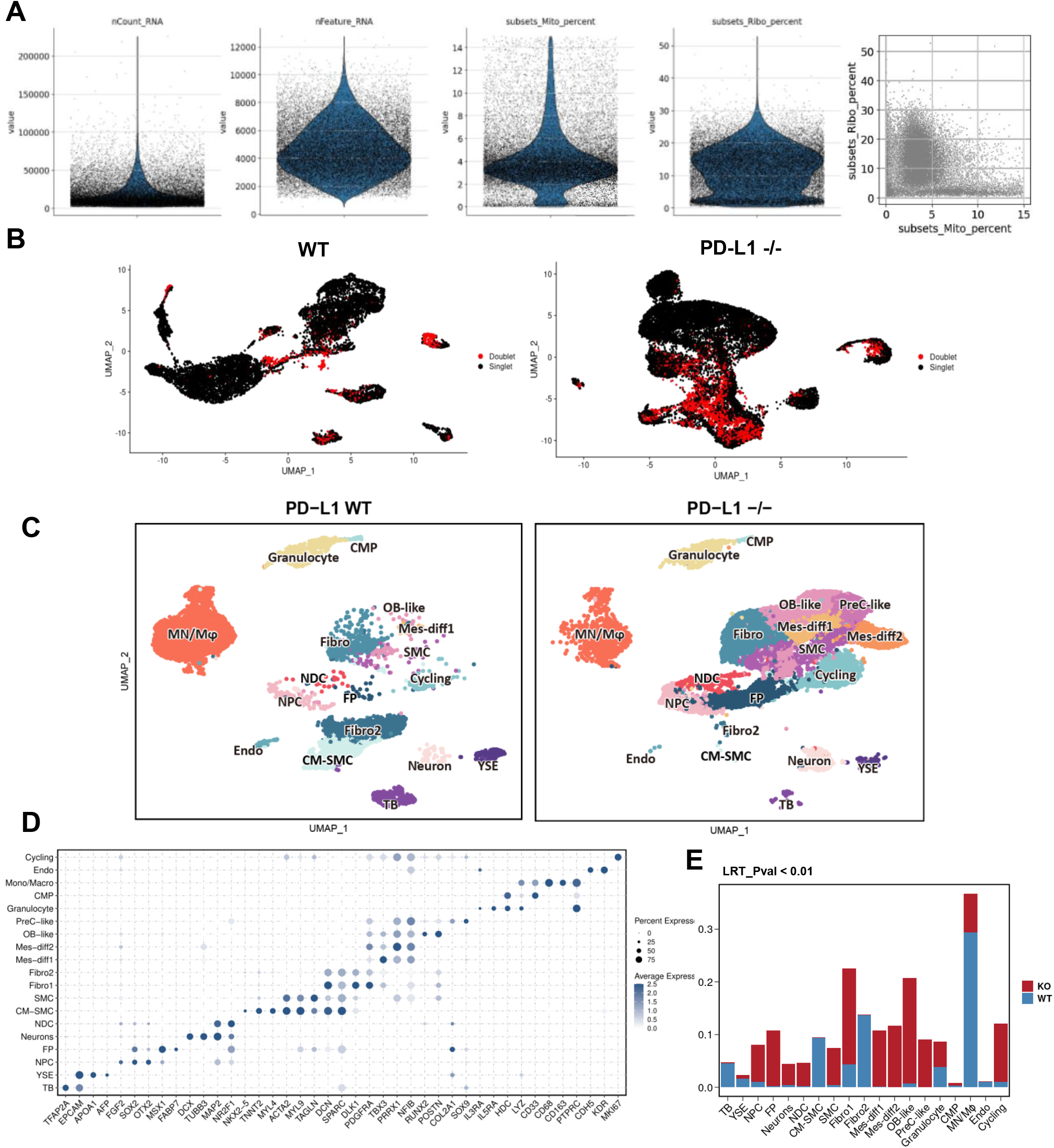
Quality control of single-cell RNA sequencing dataset and the correlation analysis with fetal yolk sac scRNA-seq dataset. **A**. Quality control of single-cell RNA sequencing dataset. Violin plots show the total number of RNA counts, total feature(gene) counts, percentage of mitochondrial gene expression, and the relative proportion of mitochondrial and ribosomal gene expression of each cell within both wild-type and *PD-L1* knockout samples. **B**. Detection of doublets and singlets in 2 samples based on DoubletFinder R package. **C**. UMAP visualization of single cells from organoids in wild-type (*WT*) and *PD-L1* knock-out (*PD-L1*^*-/-*^). Each dot represents a single cell. In total 19 cell types were annotated and displayed in different colors. **D**. Dot plot showing the expression of marker genes used for cell type annotation. The dot size represents the percentage of cells having the corresponding gene expression and the color intensity indicates the average expression of each gene. **E**. Stacked-bar plot showing the change of cell-type proportion of each between WT and KO samples. Statistical significance test (LRT_Pvalue) is similarly used in Figure 2D.

**Supplementary Figure 5.**
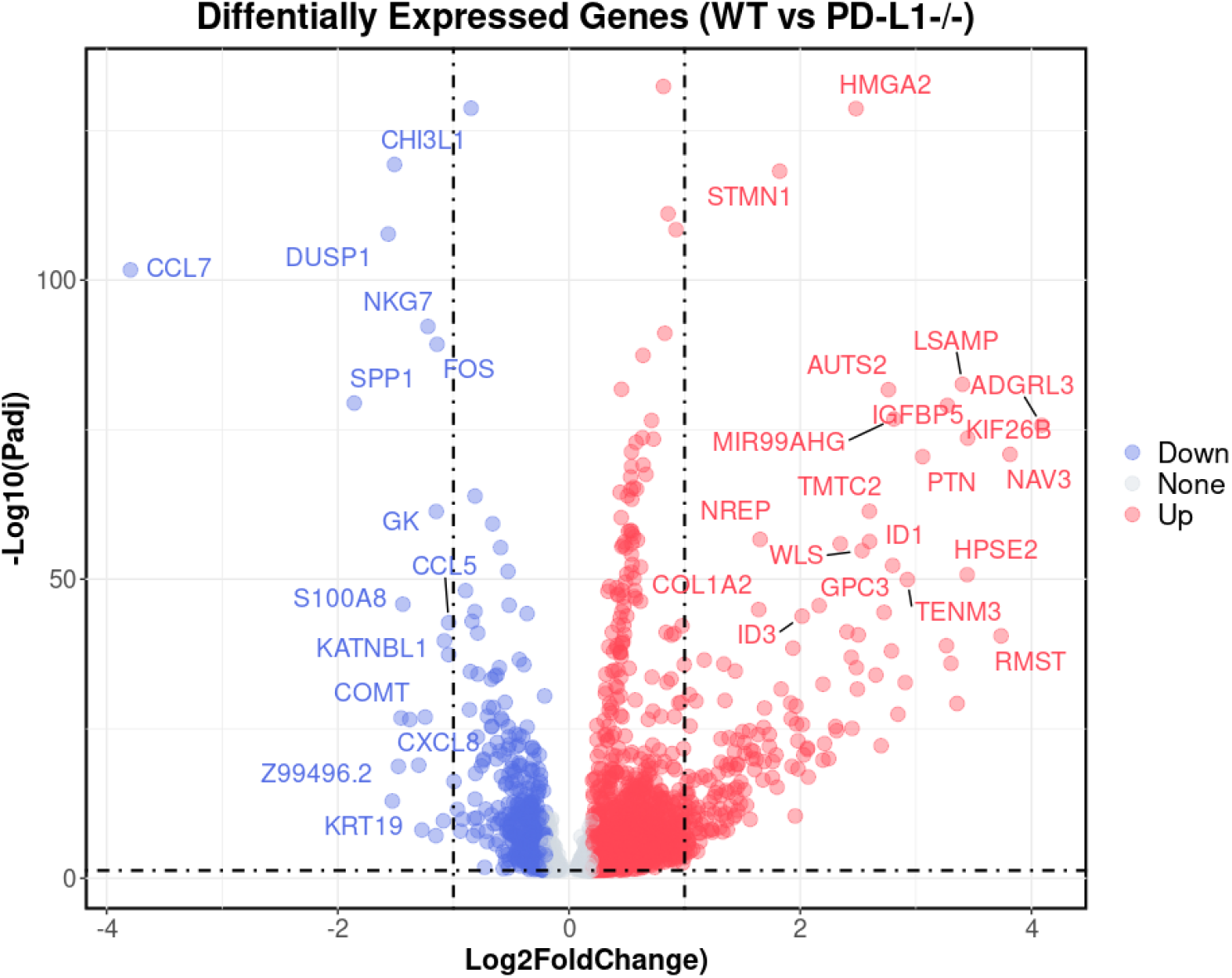
Differentially expressed genes in *WT* vs. *PD-L1 KO* MC/Mϕ. Volcano Plot showing the differentially expressed gene (DEGs) calculated by Seurat R package built-in “bimod” method, between macrophages in wild-type (*WT*) and *PD-L1* knockout (*PD-L1*^*-/-*^). DEGs (in blue/red color) are defined by q-value (calculated by “Benjamini-Hochberg” method in default) less than 0.05, and log2-fold change in an absolute value larger than 0.2. Gene symbols labeled are DEGs with log2-fold change in an absolute value larger than 1. DEGs in blue or red color are genes that are down-regulated or up-regulated upon *PD-L1* knockout, separately.

**Supplementary Figure 6.**
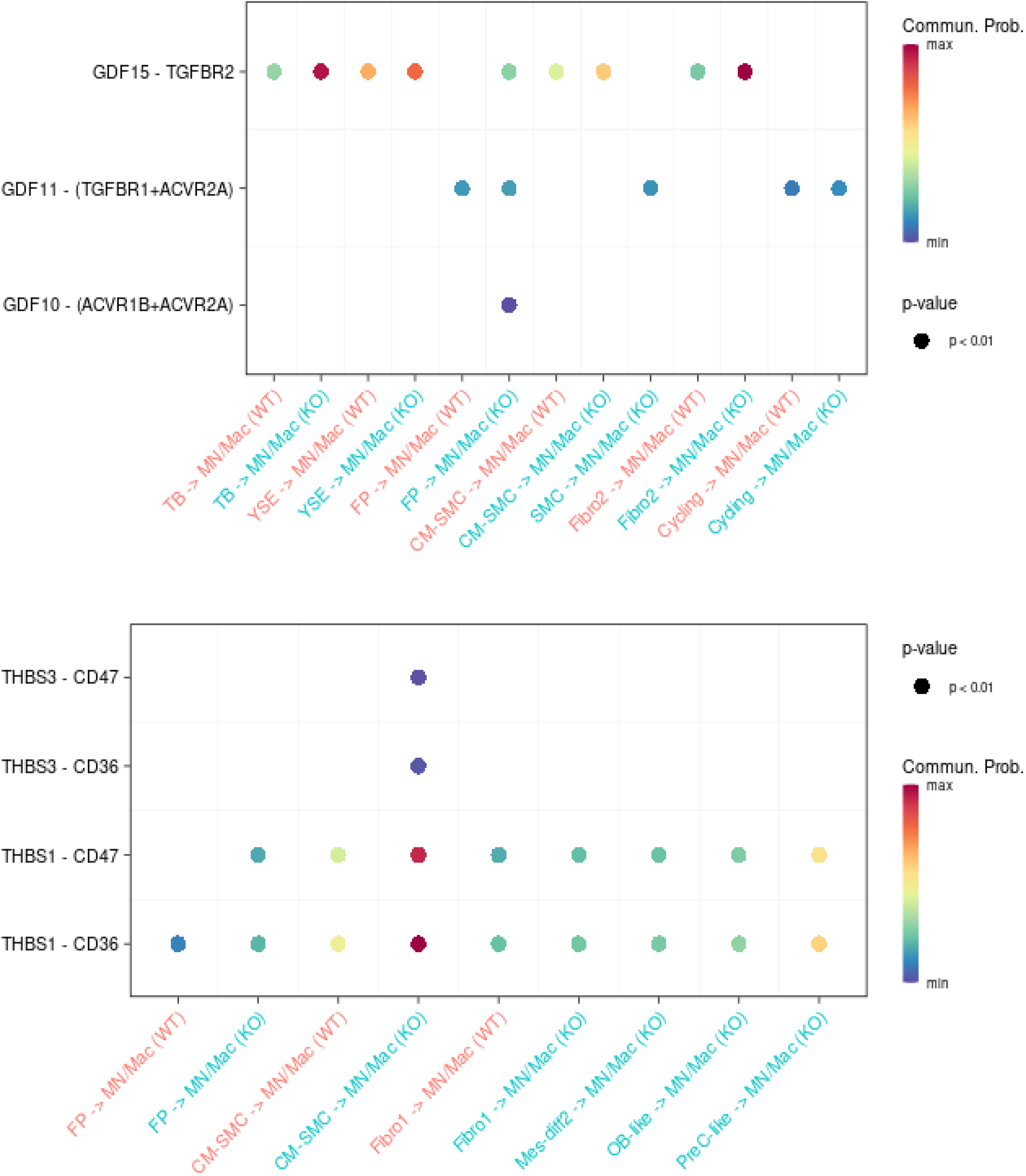
Significant interactions of GDF and THBS signals increased in macrophages upon *PD-L1* knockout. Dot-plot showing the relative significant interactions (ligand-receptor pair) of MC/Mϕ for GDF and THBS signaling pathways based on their average expression. The dot color and size represent the communication probability and p-values calculated by CellChat.

**Supplementary Figure 7.**
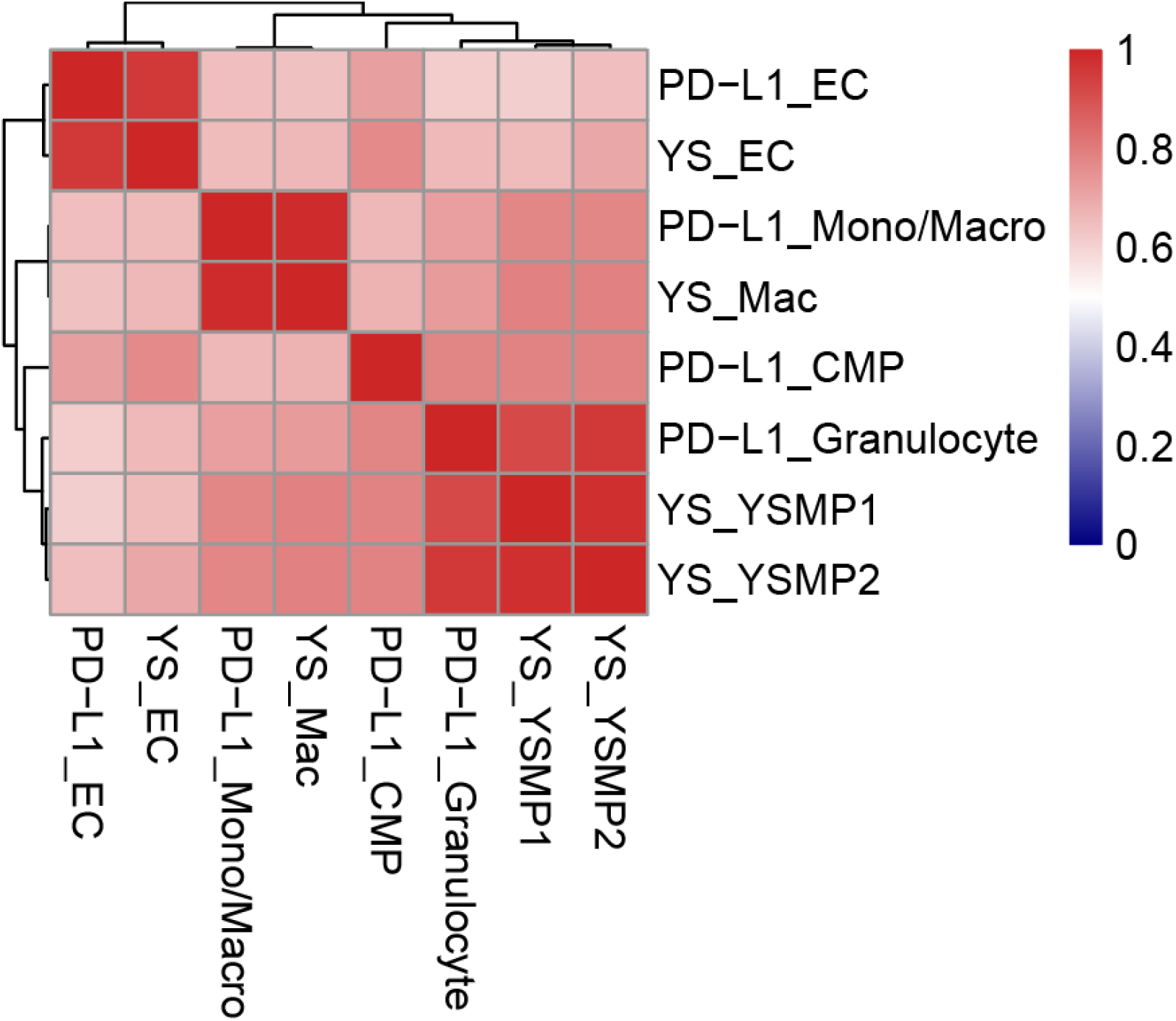
Correlation analysis with fetal yolk sac scRNA-seq dataset. Transcriptomic data were used for correlation analysis between human in vivo fetal yolk sac (YS) hematopoietic clusters (YS_EC, endothelial cells; YS_Mac, macrophages; YS_YSMP1,2, myeloid-biased progenitors) and PD-L1 WT hematopoietic cell populations (PD-L1_EC, endothelial cells; MC/Mϕ, monocyte/macrophages; CMP, common myeloid progenitors; Granulocytes).

